# Identification of the ventral occipital visual field maps in the human brain

**DOI:** 10.1101/035980

**Authors:** Jonathan Winawer, Nathan Witthoft

## Abstract

The location and topography of the first three visual field maps in the human brain, V1-V3, are well agreed upon and routinely measured across most laboratories. The position of 4^th^ visual field map, ‘hV4’, is identified with less consistency in the neuroimaging literature. Using magnetic resonance imaging (MRI) data, we describe landmarks to help identify the position and borders of hV4. The data consist of anatomical images, visualized as cortical meshes to highlight the sulcal and gyral patterns, and functional data obtained from retinotopic mapping experiments, visualized as eccentricity and angle maps on the cortical surface.

Several features of the functional and anatomical data can be found across nearly all subjects and are helpful for identifying the location and extent of the hV4 map. The medial border of hV4 is shared with the posterior, ventral portion of V3, and is marked by a retinotopic representation of the upper vertical meridian. The anterior border of hV4 is shared with the VO-1 map, and falls on a retinotopic representation of the peripheral visual field, usually coincident with the posterior transverse collateral sulcus. The ventro-lateral edge of the map typically falls on the inferior occipital gyrus, where functional MRI artifacts often obscure the retinotopic data. Finally, we demonstrate the continuity of retinotopic parameters between hV4 and its neighbors; hV4 and V3v contain iso-eccentricity lines in register, whereas hV4 and VO-1 contain iso-polar angle lines in register.

Together, the multiple constraints allow for a consistent identification of the hV4 map across most human subjects.

## INTRODUCTION

The human brain contains well over a dozen visual field maps^1-3^. Identification of these maps has been a major success in the history of visual neuroscience. Because researchers can identify the same brain region across multiple measurements and diverse populations, scientific findings can be aggregated across studies to arrive at a better understanding of human brain function. The success of such aggregation, however, is limited by the accuracy with which a given region can be identified across individuals. There is little to no doubt about the position and borders of several maps. Primary visual cortex (V1) always lies on the Calcarine sulcus^4-7^, and its borders can be identified based on data from a single fMRI scanning session with high precision^8,9^. V2 and V3 can also be identified quite accurately and routinely, and in fact the retinotopy parameters of all three maps can be reasonably estimated from the anatomy alone^10,11^.

In contrast, the fourth visual field map has proven more difficult to characterize, with considerably less consistency in map definition across laboratories, compared to V1-V3^12-20^. There are several reasons. Compared to V1-V3, the fMRI signals in hV4 are less reliably driven by simple contrast patterns^21^, the homology with animal models is less certain^22^, imaging artifacts can affect some parts of the map in many subjects^14^, and anatomical landmarks are less frequently used in map delineation. Recent work, however, has examined the hV4 map in some detail^14,19^, suggesting that one can identify hV4 and its neighbors (Figure 1) with reasonably good consistency across individuals.

**Figure 1.**
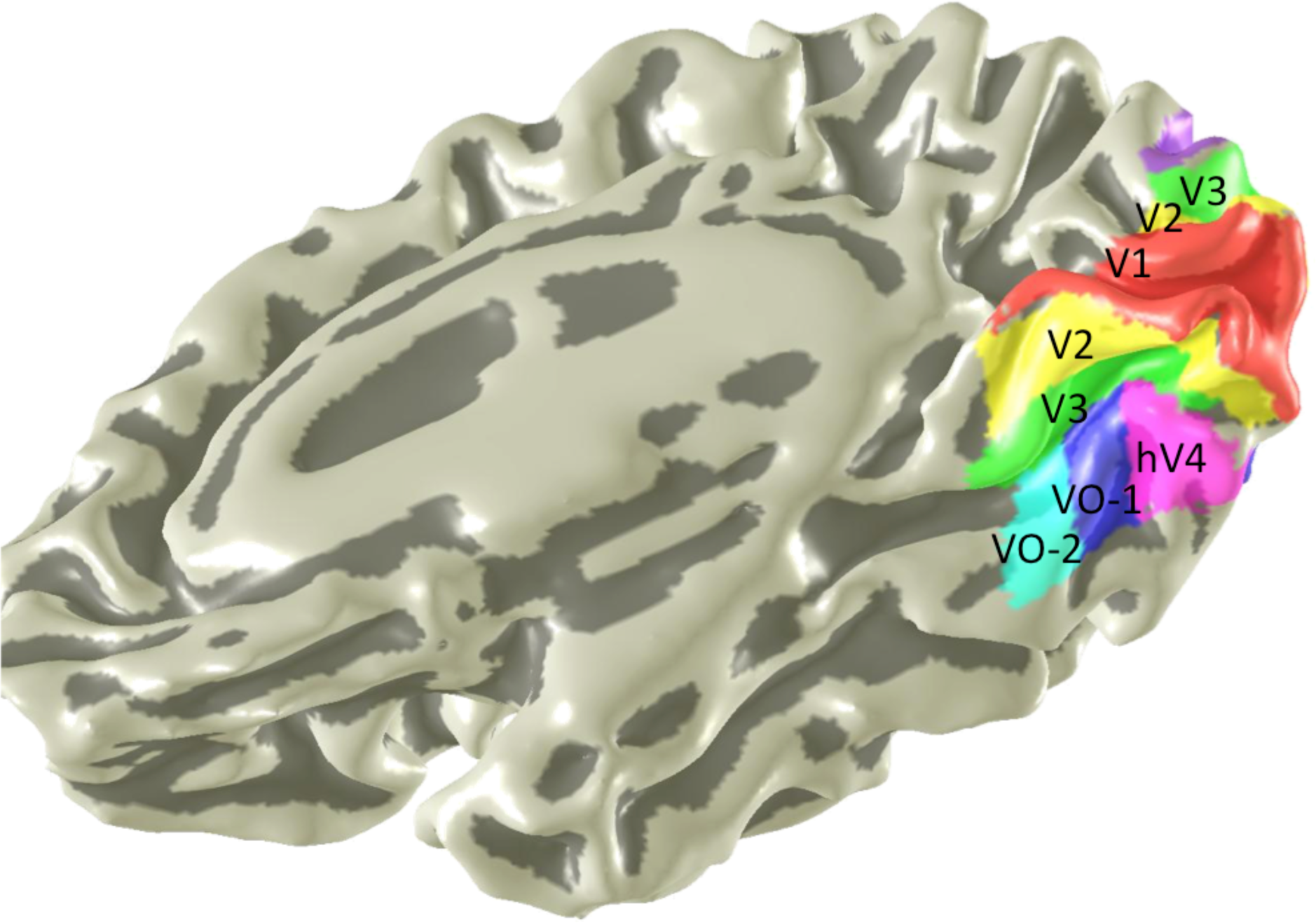
Location of ventral occipital visual field maps. The locations of 6 medial and ventral visual field maps are shown on a rendering of a subject’s right hemisphere. The mesh underlay is a slightly smoothed rendering of the cortical surface. Sulci are indicated by dark gray and gyri by light gray. Adapted from ^1^.

In our view, successful map identification of the hV4 maps is aided by combining several sources of data. First, we consider the anatomy, especially the pattern of sulci and gyri. Second we use the angle map and eccentricity map derived from retinotopy measurements. Third, we use a metric of signal quality such as mean BOLD signal (or signal to noise ratio, or the quality of the retinotopic fit) to identify possible imperfections in the measurements that should be ignored in identifying the map. In the simplest scenarios, all constraints agree and the map is easily identifiable. In some cases, the measurements do not fully agree but we can nonetheless identify the general location of hV4 and at least one or more of the map boundaries based on the mutual constraints.

The protocol for identifying the hV4 retinotopic map is described in the following text and in a step-by-step video description (Supplementary Movie 1).

## PROTOCOL

Our procedure for identifying visual field maps involves two kinds of MRI data and several processing steps. The required data are an anatomical image of the subject’s brain (structural MRI) and functional data from a retinotopic mapping experiment. Typically, all data can be collected in less than one hour of scan time on a 3 Tesla scanner using a standard gradient echo pulse sequence for functional data and T1-weighted images for anatomical data. The analysis steps include segmenting gray and white matter to generate a cortical mesh, identifying the sulcal and gyral patterns in ventral occipito-temporal cortex, extracting the eccentricity and angle data from the retinotopic experiments, and then using these measurements to guide the marking of visual field boundaries.

There is considerable variability in the specific procedures used by different research groups for acquisition of functional and anatomical MRI data and for the derivation of retinotopy parameters from the fMRI scans. Any of the retinotopy methods in common usage is suitable; the only requirements are that one can visualize eccentricity and angle data on a cortical mesh. The methods to produce the images shown in this paper are described in more detail elsewhere^19^. We summarize these methods briefly and then focus on the specific steps for identifying the hV4 map and its neighbors.

### 1. Data acquisition

#### 1.1 Anatomical data

Anatomical data are needed in order to render retinotopic data on surface representations. For subjects 1-3 depicted in the video presentation, 2-4 whole brain SPGR T1-weighted MRI images were acquired on a 3T GE scanner at 1 mm isotopic resolution using an 8-channel whole-brain coil. The multiple images for each subject were aligned and averaged. Averaging multiple anatomical images is desirable as it increases the contrast of the boundary between the grey and white matter and therefore aids segmentation and the creation of an accurate cortical surface.

#### 1.2 Functional data

Functional data were acquired as T2*-weighted gradient echo images with either a spiral ^23^ or rectilinear (EPI) trajectory through k-space, with 2.5 mm isotropic voxels and a TR of either 1.5 s or 2 s. (For details see ref ^19^.)

### 2. Stimuli

Retinotopy was measured with stimuli designed either for travelling wave analysis^24^ or population receptive field modeling^25,26^. For traveling wave analysis, alternate scans contained contrast patterns viewed through expanding ring apertures or clockwise rotating wedges with a visual field extent of 14 degrees from fixation. (For details see ref ^19^.) For population receptive field modeling, stimuli were contrast patterns viewed though moving bar apertures that slowly swept across the visual field, with a maximum eccentricity of 6 degrees from fixation. (For details see refs ^14,25^)

### 3. Functional MRI analysis

#### 3.1 Preprocessing

Several standard fMRI preprocessing steps were used prior to retinotopic analysis, including slice timing correction, motion correction, and high pass temporal filtering with a 1/20 Hz cutoff.

#### 3.2 Travelling wave analysis

Data from traveling wave scans (wedges and rings) were analyzed voxel by voxel by applying the Fourier transform to the time series. The phase at the stimulus frequency measures the polar angle or eccentricity of the stimulus that most effectively drives the BOLD response in that voxel. The coherence of the signal at the stimulus frequency indicates the goodness of fit of the response to the periodic design. Coherence is defined as the power at the stimulus frequency divided by the power across all frequencies. For details see ref ^27^.

#### 3.3 Population receptive fields

A population Receptive Field (pRF) model was solved for each voxel using methods described previously^25^. In brief, each pRF was defined as a 2-D isotropic Gaussian, parameterized by its center (x, y), size (sd of Gaussian), and amplitude (scale factor). The model prediction is the dot product of the pRF and the stimulus aperture, convolved with a hemodynamic response function. The model was fit through a minimization of least squared error between the predicted and observed time series using a coarse-to-fine approach. The pRF center can be converted to polar coordinates to yield an angle and eccentricity map, similar to that obtained from traveling wave analysis. For the purposes of retinotopic mapping in this paper, other pRF parameters were not used (size and amplitude).

### 4. Segmentation and visualization

Because the retinotopic maps in visual cortex are organized on the 2D surface, not the 3D volume, map data is best visualized on a surface representation, typically rendered as an inflated (smoothed) or un-inflated 3D surface mesh, or as a flattened (2D) surface ^28-30^. The data used for the surface rendering is usually the border of the gray and white matter, derived from a segmentation procedure that labels anatomical voxels from the T1 image as gray or white matter, and finds then finds the boundary that separates the two tissues. For the data shown in this paper, we segment the gray and white matter using the Freesurfer auto-segmentation tools^31^ followed by manual correction using ITK-GRAY software (http://white.stanford.edu/software/, modified from ITK-SNAP http://www.itksnap.org)^32^. Functional data were then aligned to the whole brain anatomical data and rendered on a smoothed surface using custom software (vistasoft; https://github.com/vistalab/vistasoft; http://vistalab.stanford.edu/software/).

### 5. Anatomical landmarks

We find it helpful to identify several sulci and gyri before drawing the retinotopic maps, and follow the naming conventions used by Duvernoy^33^. The sulci and gyri labeled in Figure 2 can all be seen in post-mortem brains in Duvernoy’s text (e.g., his figure 17). Their locations are described briefly below.

**Figure 2.**
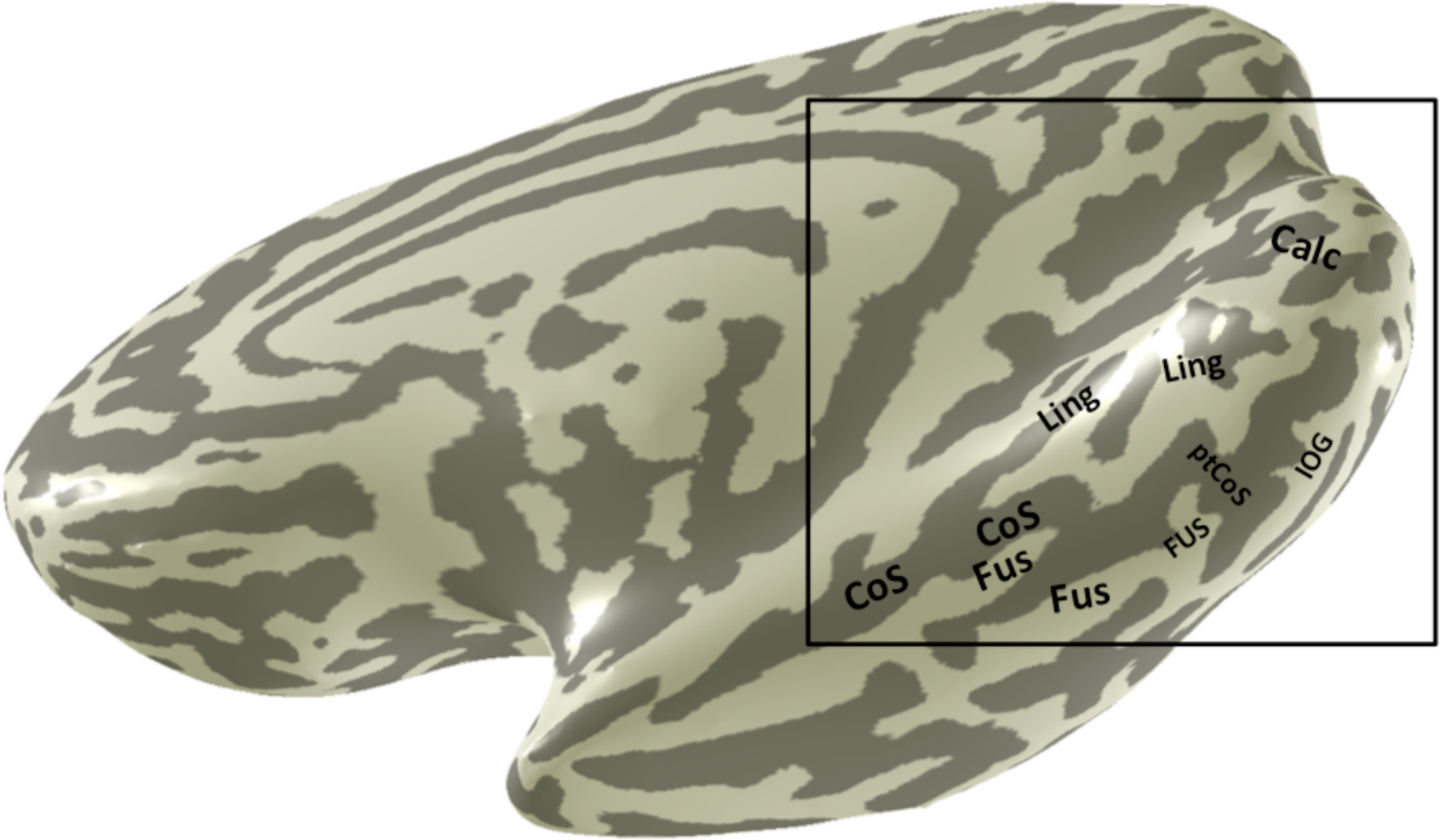
Anatomical landmarks in ventral occipitotemporal cortex. Several sulci and gyri are labeled on the ventral and medial surface of a highly smoothed right hemisphere. These sulci and gyri provide useful landmarks in the identification of the visual field maps. Calc: Calcarine sulcus; Ling: Lingual sulcus/gyrus; CoS: Collateral sulcus; Fus: Fusiform gyrus; ptCoS: Posterior transverse collateral sulcus; IOG: Inferior occipital gyrus.

#### 5.1 Calcarine sulcus

The most useful landmark is the calcarine sulcus (sometimes called the calcarine fissure), which locates primary visual cortex (V1). It is the large sulcus in medial occipital cortex separating the lingual gyrus from the cuneus. It can be found unambiguously in every subject and is always the location of V1. It runs on an anterior-posterior axis, with the anterior end marked by the parietal occipital fissure, and the posterior end approximately at the occipital pole, though there is variability in the posterior extent, with the sulcus wrapping around to the lateral surface in some subjects, and terminating on the inferior surface in others^34,35^.

#### 5.2 Lingual gyrus and lingual sulcus

The lingual gyrus is inferior to the calcarine sulcus and is the location of the ventral portions of V2 and V3. Its inferior boundary is the collateral sulcus. It stretches approximately from the posterior pole to the parahippocampal gyrus. The lingual gyrus contains one or more sulci together considered the lingual sulcus.

#### 5.3 Collateral sulcus

The collateral sulcus separates the lingual and parahippocampal gyri on the one side from the fusiform gyrus on the other. Its posterior end is usually denoted by a transverse portion of the sulcus, called the posterior transverse collateral sulcus (ptCoS). The ptCoS is an important landmark because it usually marks the division between the hV4 map and the VO-1 map^19^.

#### 5.4 Fusiform gyrus

The fusiform gyrus is inferior to the lingual and parahippocampal gyrus, and is bounded medially by the collateral sulcus. The foveal representation of the ventral-occipital maps (VO-1/2) lies in the posterior portion of the fusiform gyrus.

#### 5.5 Inferior occipital gyrus

The inferior occipital gyrus lies posterior to the fusiform gyrus and collateral sulcus and inferior to the lingual gyrus. It is generally separated from the posterior portion of the posterior fusiform gyrus by the posterior transverse collateral sulcus. The lateral portion of the hV4 map often lies here. This gyrus is typically in close proximity to the transverse venous sinus, which can lead to fMRI artifacts that obscure this portion of the hV4 map^14^.

### 6. Drawing the maps

#### 6.1 V1-V3 maps

Identification of the first three visual field maps in humans was one of the first accomplishments in visual neuroscience following the advent of fMRI^24,27,36^. Delineation of these map boundaries is now routine, although tracing the maps through the foveal confluence can be difficult in the absence of methods developed specifically for this purpose^9^. In brief, the V1 map is centered on the calcarine sulcus, and its borders with V2 lie on retinotopic representations of the vertical meridian. The peripheral boundary is usually not identified with functional MRI because the field of view of that can be achieved during scanning is less than the field of view represented in the maps. Hence the peripheral boundary of a region of interest is usually chosen as the most anterior region of the maps where the signal is good, according to a threshold in coherence (traveling wave analysis) or variance explained (pRF analysis). If there are clear reversals in the angle map all the way back to the occipital pole, then it may be possible to trace V2 and V3 in concentric ‘V’ shapes around V1; otherwise the most foveal portion of the maps is typically labeled as a non-specific ‘foveal confluence’, and each map is traced as far into the fovea as the resolution in the angle maps allow, usually one or two degrees from fixation, such that the two arms of the ‘V’ are not connected, and labeled as the dorsal and ventral portions of the map The dorsal and ventral V2/V3 borders are marked by a representation of the horizontal meridian in the angle map.

#### 6.2 The hV4 map

The hV4 map lies on the ventral surface of the occipital lobe and borders at least two other visual field maps, the ventral part of V3 (V3v) and VO-1.

##### 6.2.1. The anterior boundary: hV4 / VO-1

The boundary between hV4 and VO-1 can be identified by both anatomical and retinotopic data. The retinotopic feature that defines this border is a reversal in the eccentricity map (Figure 3). This peripheral eccentricity band dividing hV4 from VO-1 usually coincides with a particular anatomical feature, the posterior transverse collateral sulcus (ptCoS)^19^.

**Figure 3.**
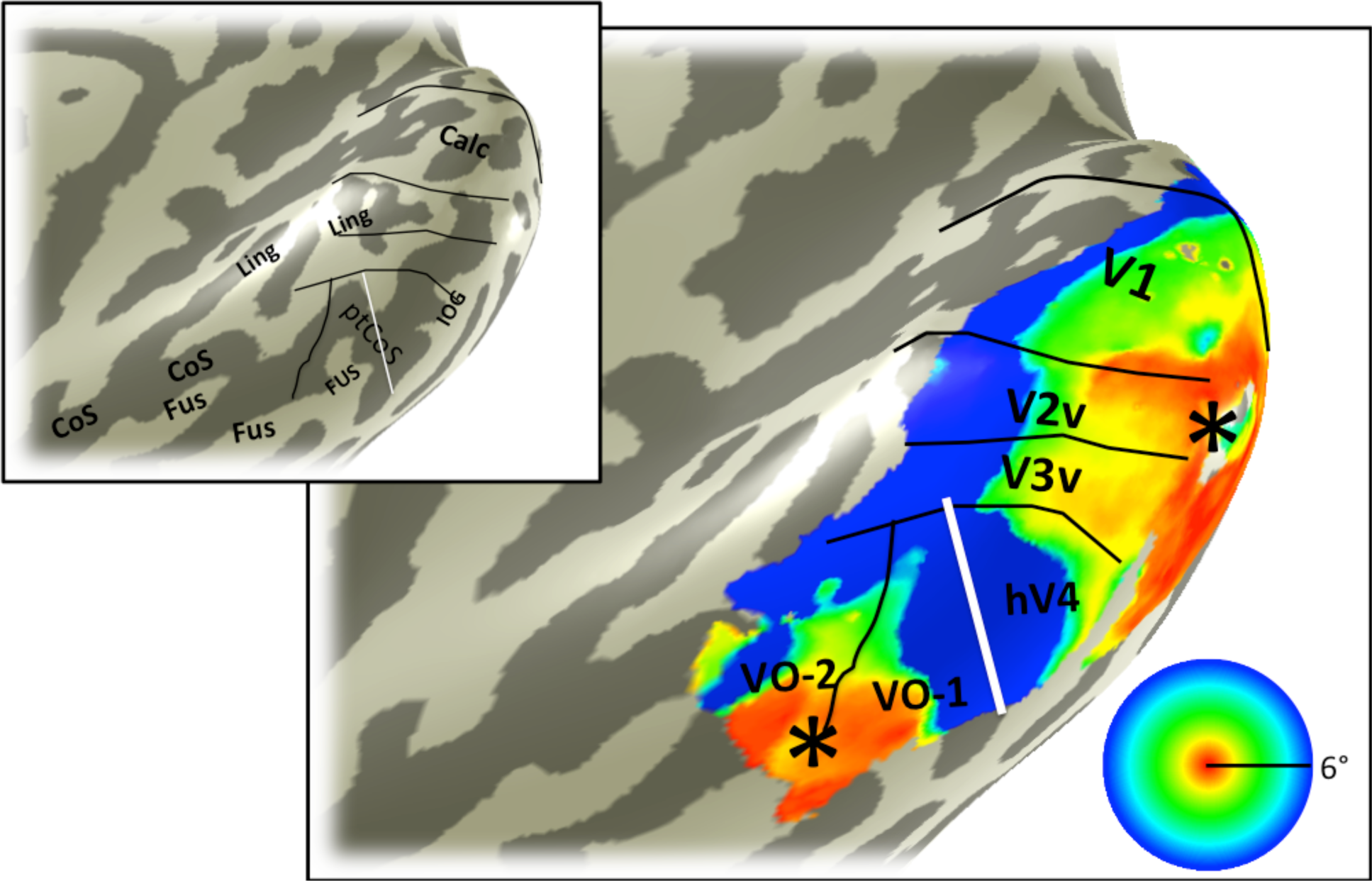
Eccentricity map. The color overlay on the smoothed cortical mesh in the main panel shows the stimulus eccentricity that most effectively drives each cortical location. The eccentricity was derived from solving a pRF model, and the stimulus extent was limited to 6 degrees. The asterisk indicates the confluent foveal representation. Black and white lines mark the boundaries between visual field maps. The white line divides hV4 from VO-1 and is the only boundary line in this figure that is derivable from the eccentricity map. This line coincides with the ptCoS (inset) and with the eccentricity reversal (blue in color overlay). Hence the white line also divides the ventral occipital maps into two clusters, one that includes V1-hV4, and one that includes VO-1/2^16^. Data are limited to voxels in which the pRF prediction accounted for at least 10% of the variance explained in the time series and to a posterior mask that includes the 6 labeled visual field maps: V1, V2 and V3 ventral, hV4, VO-1, and VO-2. The inset is the same cortical mesh with labels indicating the major sulcal and gyral patterns, as in figure 2.

##### 6.2.2. The medial boundary: hV4/V3v

One of the borders of hV4 is shared with the ventral portion of V3 (V3v). This border is defined by an upper vertical meridian polar angle reversal (Figure 4). Because the V1-V3 maps are typically well defined by the retinotopic data, it is usually also clear where this boundary is found.

**Figure 4.**
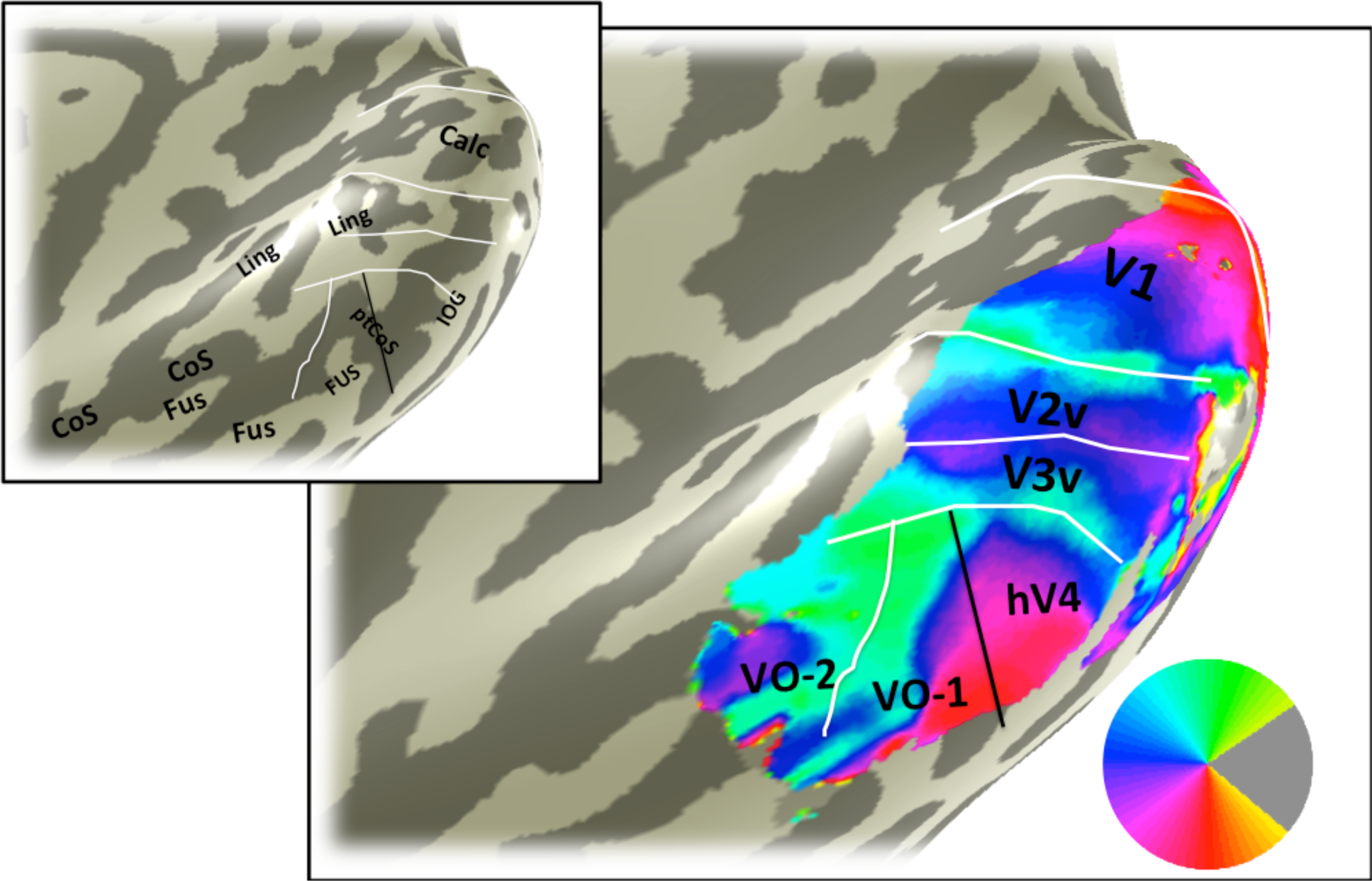
Angle map. The color overlay in the main panel indicates the angle in the visual field that most effectively drives responses in each cortical location. The visual field map boundaries in white can be identified from the angle map. Otherwise as figure 3.

##### 6.2.3. The ventral/lateral boundary

The ventral/lateral boundary of hV4 is more difficult to identify than the other boundaries because there is no unambiguous feature of the retinotopic data to define it. As described in 6.2.1 and 6.2.2, we know what is on the other side of two of the hV4 boundaries, making those boundaries well defined: The anterior boundary is defined by an eccentricity reversal and is shared with VO-1, and the medial boundary is defined by a polar angle reversal and is shared with V3v. In contrast, there is not a well-established map or retinotopic feature abutting the ventral side of hV4. Furthermore, there is often signal dropout in the fMRI measure on the ventral / lateral aspect of the hV4 map due to the transverse venous sinus, which typically runs near the inferior occipital gyrus^14^. In some cases, the signal dropout defines the most ventral/lateral extent of the map that can be identified (Figure 5; Figure 6).

**Figure 5.**
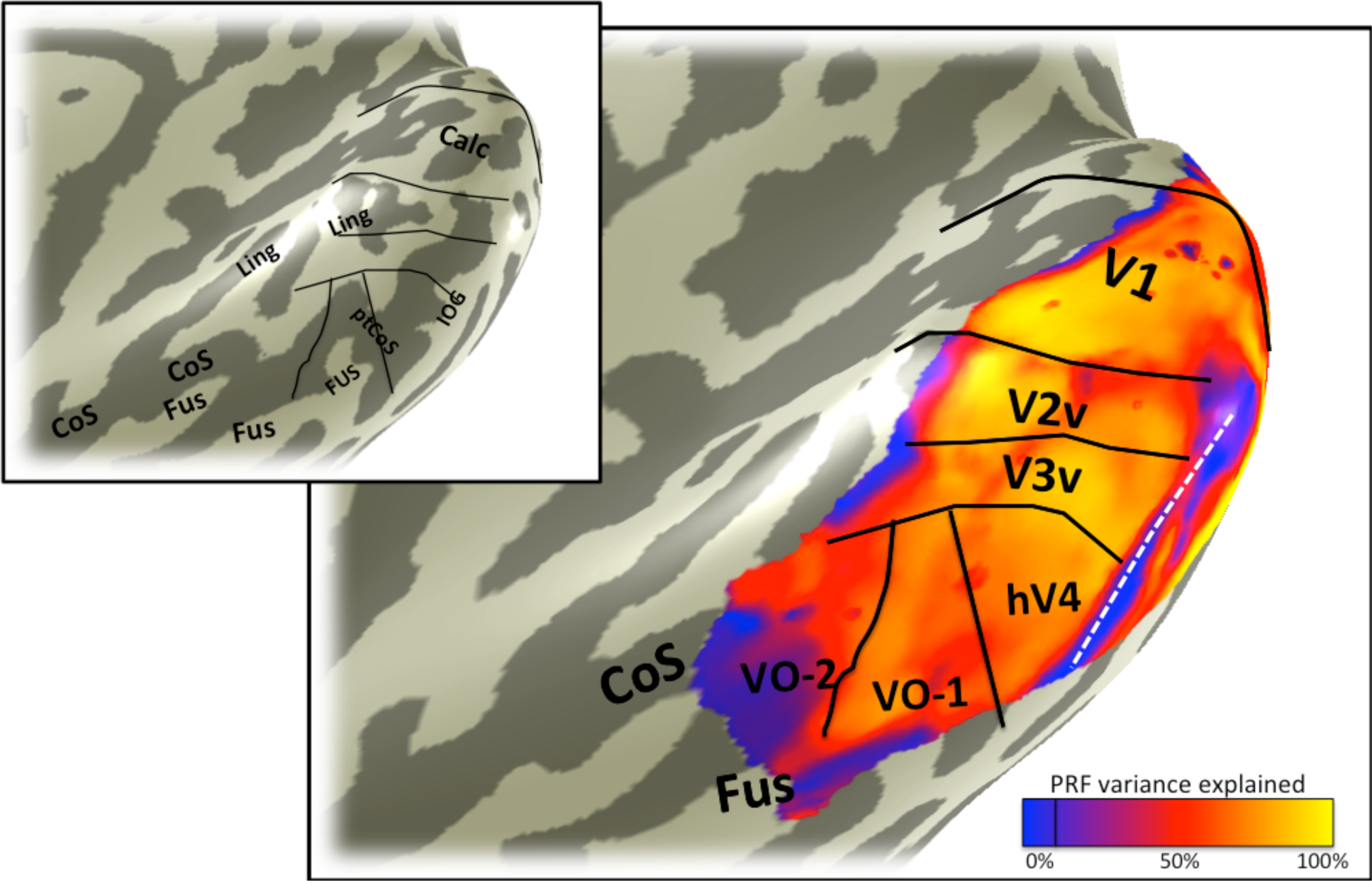
Variance explained map. The color overlay in the main panel indicates variance in the voxel time series explained by the pRF model predictions. Otherwise as figure 3. Some locations with low variance explained are likely due to fMRI artifacts, such as the region indicated by the white line, where the low variance explained is caused by dropout from the transverse sinus. Other regions with low variance explained may have visual field representations outside the stimulus extent, such as the anterior edge of V1, V2, and V3.

**Figure 6.**
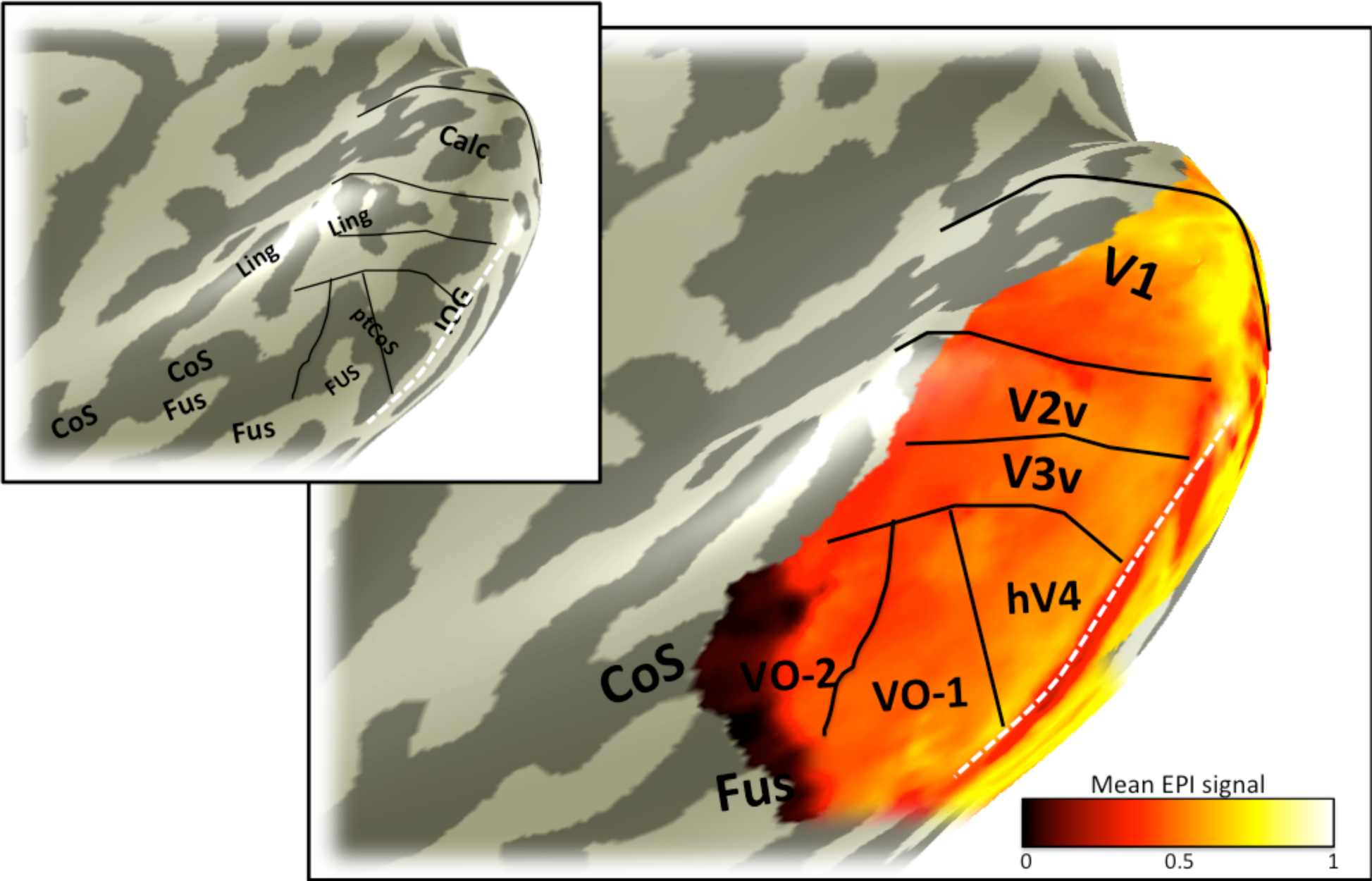
Mean signal map. The color overlay in the main panel indicates the mean fMRI signal. The signal is not uniform across cortex. Certain regions have low mean signal due scanning artifacts. The region indicated by the white line lies near the transverse venous sinus which causes signal dropout. Retinotopic data from these locations must be interpreted with caution. Otherwise as figure 3.

##### 6.2.4. Iso-eccentricity lines shared by hV4 and V3v

The general organization of the hV4 map and its neighbors can be understood by tracing iso-eccentricity and iso-angle lines. The hV4 eccentricity map is in register with the V1-V3 maps, so that iso-eccentricity lines span the hV4/V3v border (Figure 7). Unlike the V1-V3 maps, the hV4 map appears to contain little representation of the far periphery; this may be because the neurons in hV4 do not respond to stimuli in the periphery, or it may be because peripheral representation is highly compressed into a small amount of cortex, such that responses to more foveal stimuli have a much larger effect on the BOLD time course, masking the peripheral representation. Along the posterior-to-anterior axis, hV4 is therefore much shorter than V1-V3, and the most anterior eccentricity band in hV4 crossed V3v near the middle of the length of V3v.

**Figure 7.**
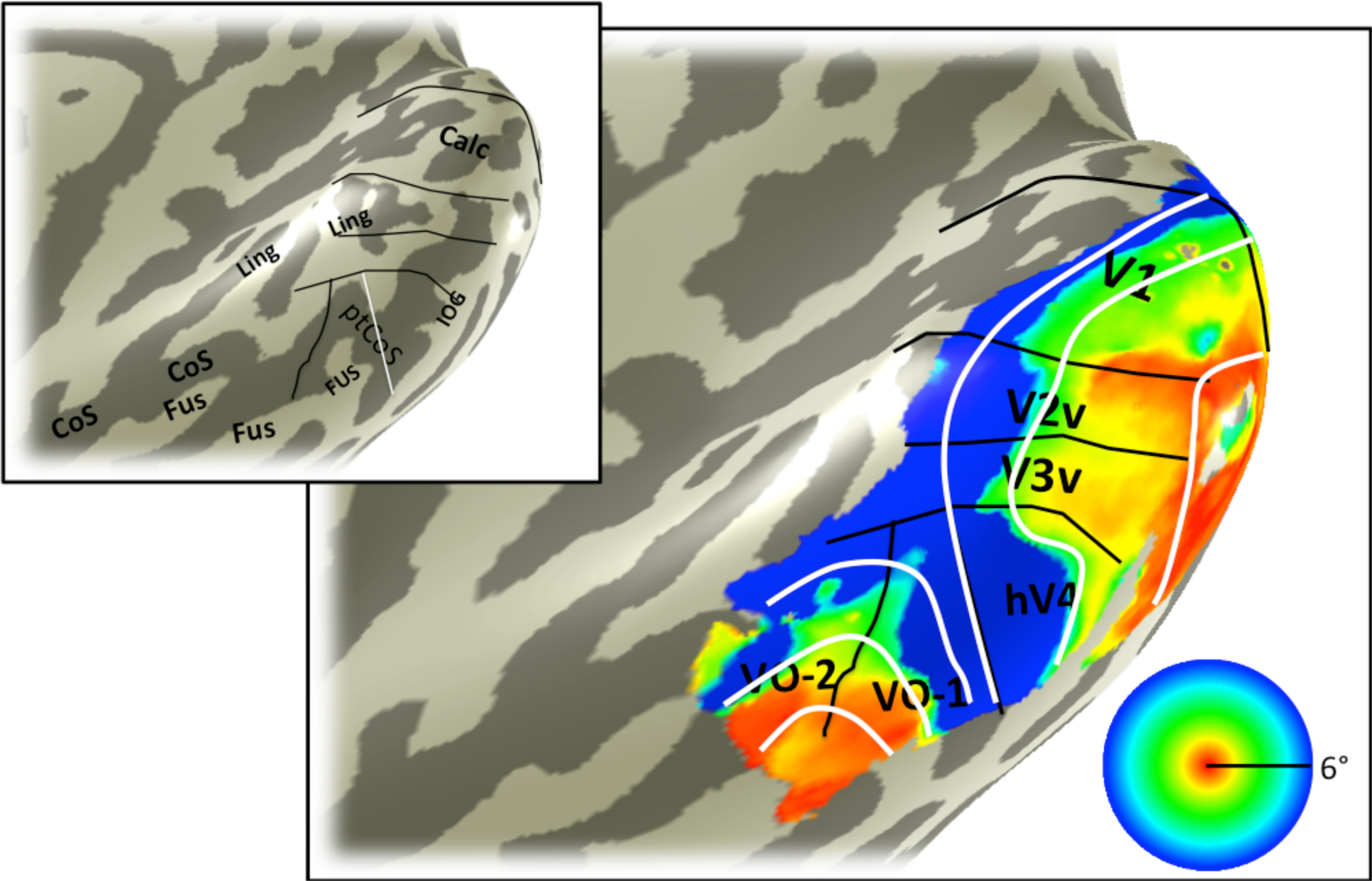
Iso-eccentricity lines. Iso-eccentricity lines in visual cortex are continuous within clusters, such as the posterior cluster containing V1, V2, V3, and hV4, and the ventral occipital cluster containing the VO-1/2 maps. The eccentricity overlay and the map boundaries are identical to those in figure 3.

##### 6.2.5. Iso-angle lines shared by hV4 and VO-1

The iso-angle lines in hV4 are continuous with the VO-1 maps and not the V3v map. The lines usually bend along the hV4/V0-1 border, with the lower meridian representation being the most ventral and the shortest iso-angle line (Figure 8).

**Figure 8.**
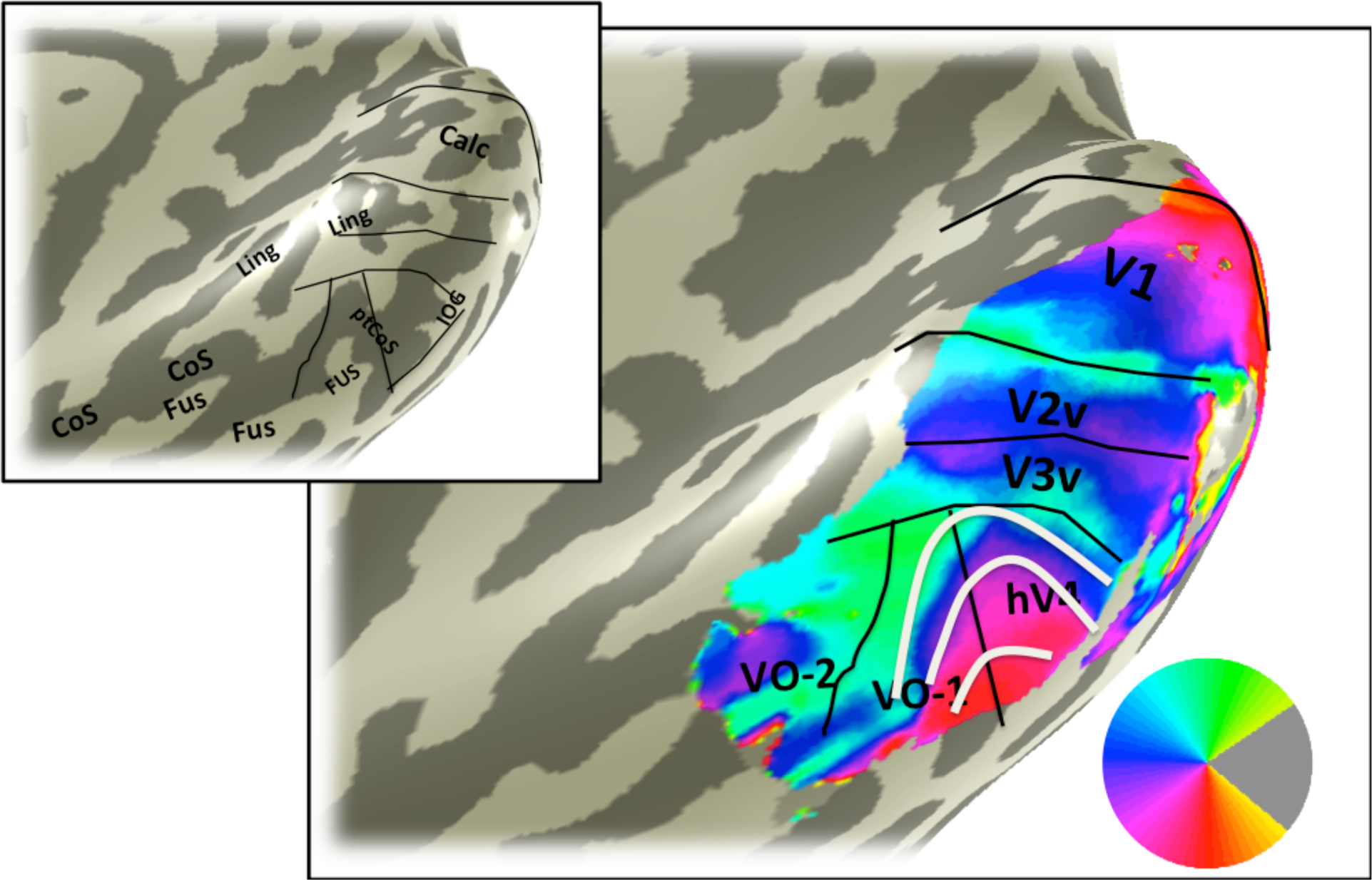
Iso-angle lines. The iso-angle lines in hV4 and VO-1, indicated by white lines, are continuous across the two map clusters. Otherwise as figure 4.

## RESULTS

The sulci and gyri associated on the ventral occipital cortical surface from a representative subject (S1) are shown in Figure 2. The most useful anatomical landmarks for locating the visual field maps are the calcarine sulcus (V1); the lingual gyrus and lingual sulcus (V2v/V3v); the posterior collateral sulcus and inferior occipital gyrus, which bound hV4; and the fusiform gyrus and collateral sulcus, where the VO maps are found.

The eccentricity map for subject S1 shows the large-scale organization of the maps (Figure 3). The key feature on the eccentricity map for identifying hV4 is the peripheral representation within the ptCoS, which marks the hV4/VO-1 boundary. The angle map (Figure 4) is used to define the V3v/hV4 boundary as well as the VO-1/VO-2 boundary.

FMRI signal quality is not uniform across the cortical surface. Some locations have poor signal due to known measurement artifacts, such as those which arise in regions near large sinuses. In subject S1, the transverse venous sinus corrupts the fMRI signal on the inferior occipital gyrus, resulting in low variance explained by the pRF model (Figure 5) and signal dropout (Figure 6) in this region.

Iso-eccentricity and iso-angle lines in subject S1 clarify the internal structure of the hV4 map and the relationship between hV4 and its neighbors. The iso-eccentricity lines are in register across hV4 and V1-V3 (Figure 7). The iso-angle lines are continuous across the hV4 / VO-1 boundary (Figure 8).

Angle maps and eccentricity maps from additional subjects show similar patterns to S1. The V1-V3, hV4 and VO-1/2 maps are shown for a representative right hemisphere (S2; Figure 9; Figure 10) and left hemisphere (S3; Figure 11; Figure 12). The large scale organization is similar for all subjects. For example, there is an eccentricity reversal dividing hV4 and VO-1 and an angle reversal dividing hV4 and V3. In all subjects, the eccentricity reversal falls on or near the ptCoS. Some details differ between subjects. For example, the foveal representation in hV4 is clear in S1 and S2 but not S3 (Figure 11), likely due to corrupted signal from draining veins. A second difference across subjects is that the upper meridian angle reversal dividing VO-1 and VO-2 is clear in S1 and S3 but not S2 (Figure 10). Nonetheless, there is sufficient regularity across subjects to identify the principal features defining the hV4 map and its neighbors, V3v and VO-1 (Figure 13).

**Figure 9:**
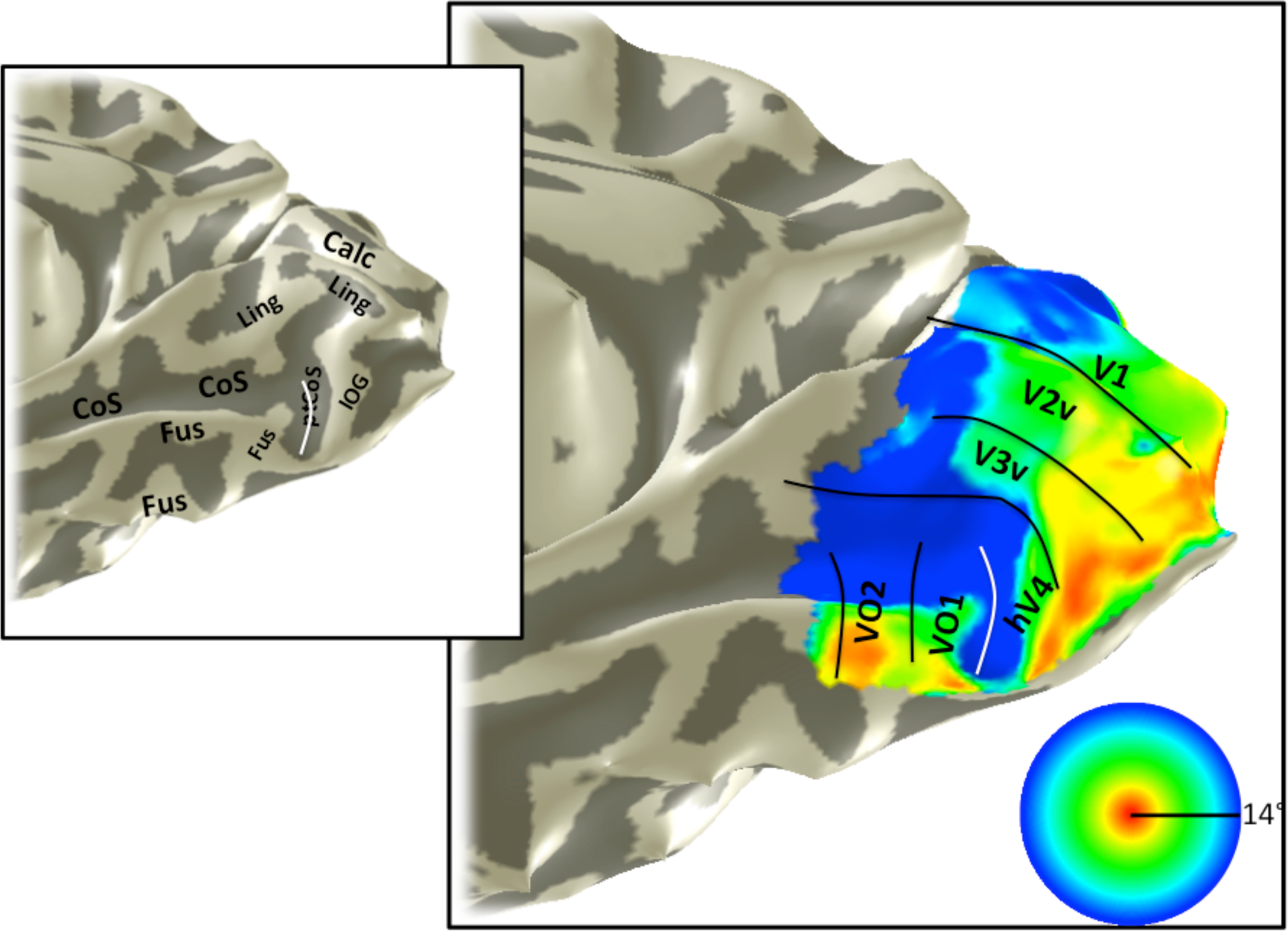
Eccentricity map, S2. An eccentricity map is shown on the partially inflated cortical surface of subject 2’s right hemisphere. The stimulus extent was 14 degrees and the data are from traveling wave analysis of expanding rings. Otherwise as figure 3.

**Figure 10:**
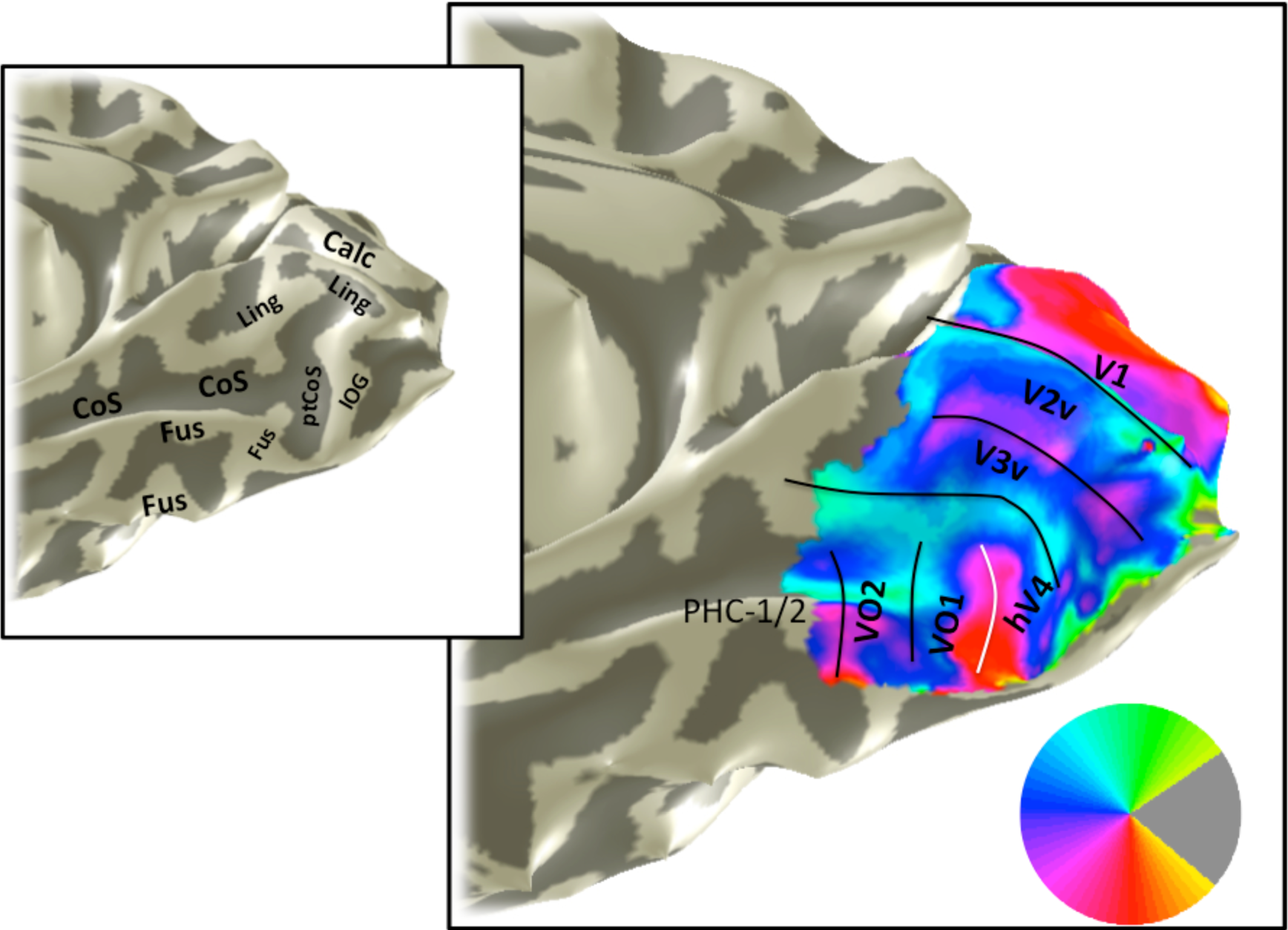
Angle map, S2. An angle map is shown for subject 2’s right hemisphere. Otherwise as figure 9 and figure 3.

**Figure 11:**
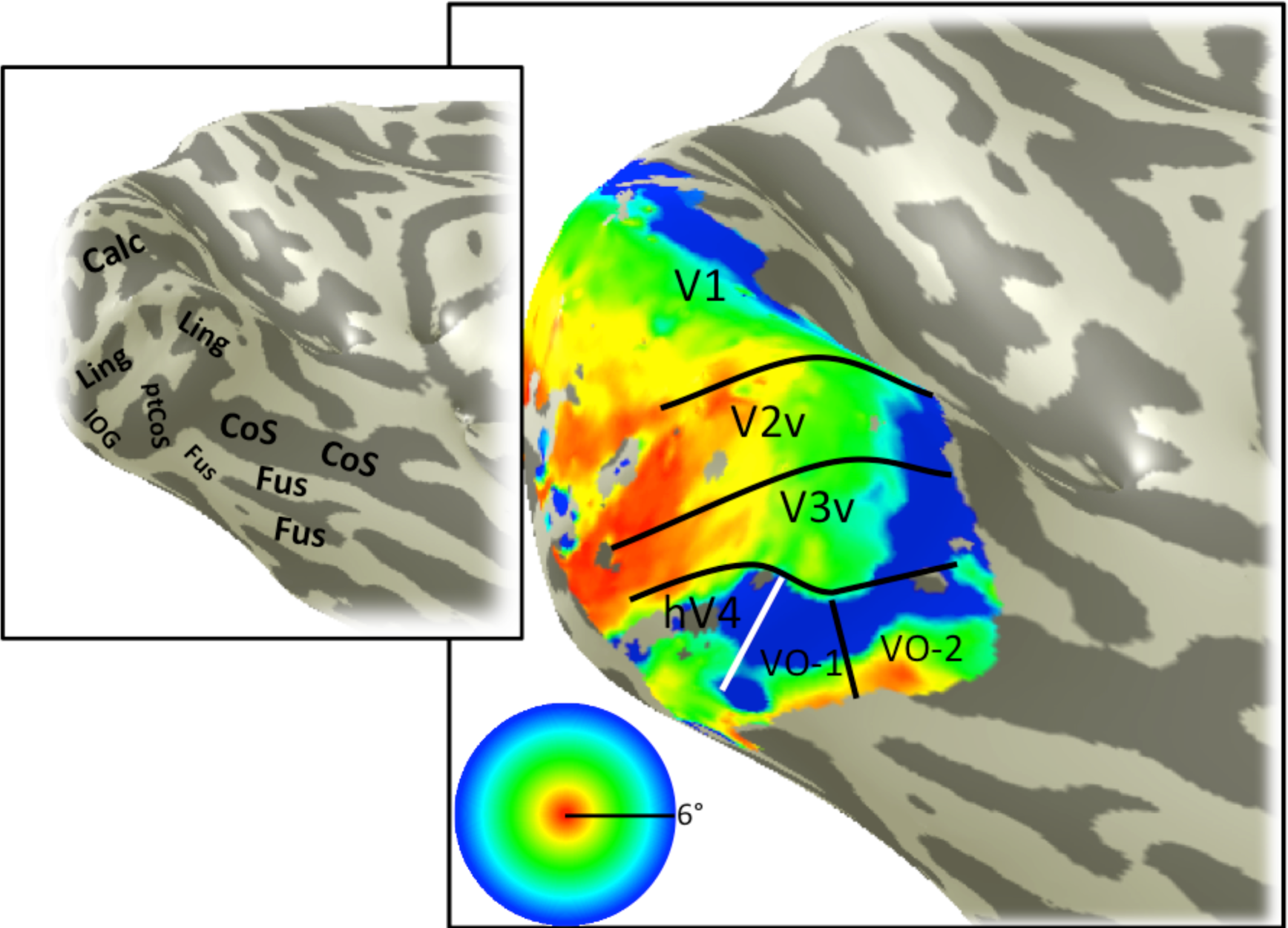
Eccentricity map, S3. An example of a left hemisphere eccentricity map derived from pRF model fitting. Otherwise as Figure 3.

**Figure 12:**
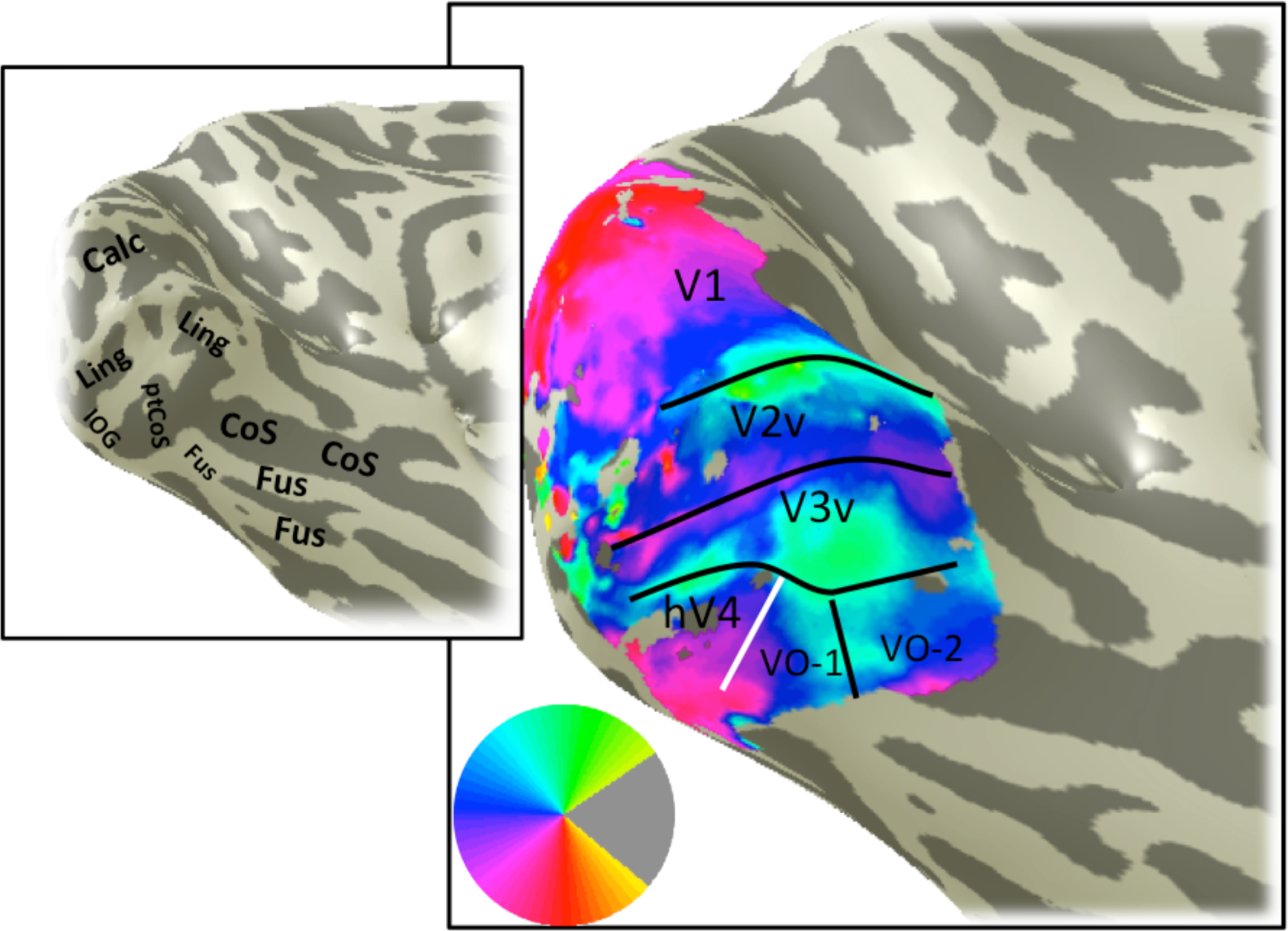
Angle map, S3. An example of a left hemisphere angle map derived from pRF model fitting. Otherwise as Figure 4.

**Figure 13:**
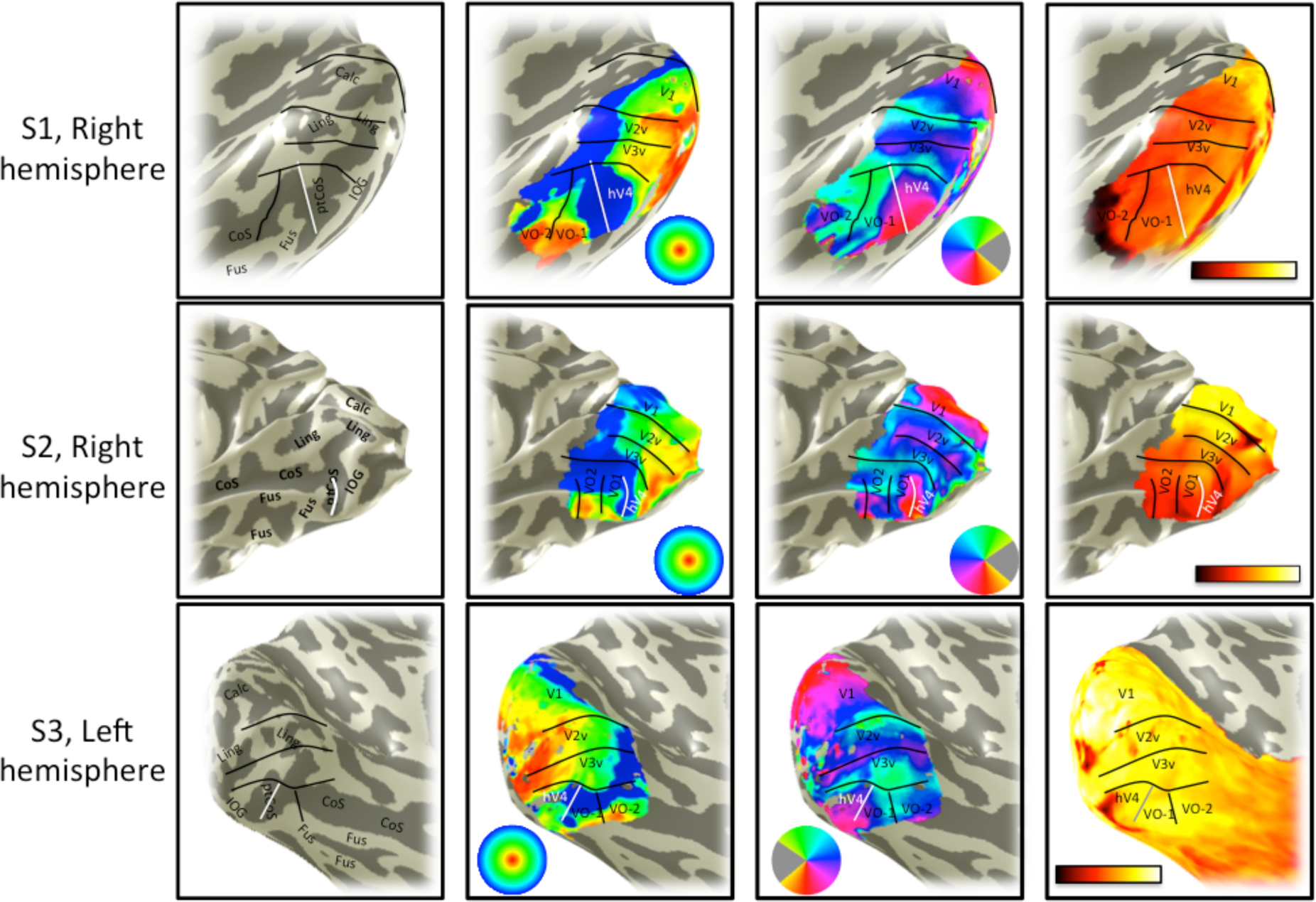
Map comparison, S1-S3. Four types of data are shown for three subjects: sulcal and gyral landmarks (column 1), eccentricity (column 2), angle (column 3), and mean fMRI signal (column 4). Comparisons of the datasets reveal regularities across subjects, such as the fact that the ptCoS is well aligned with the hV4/VO-1 boundary defined by an eccentricity reversal (white line in columns 1 and 2). There are also differences across subjects. For example the foveal representation of hV4 is less clear in S3 than in S1 and S2.

## DISCUSSION

### Regularity of maps and anatomy

Identifying visual field maps is an important component of characterizing the organization and function of visual cortex. It is important to have well-justified and reproducible methods to define the maps. In the case of the V1-V3 maps, the functional and anatomical organization is sufficiently regular and well understood that these maps can be identified using automated procedures (no human intervention)^10,11^. The hV4 and VO maps are not yet included in automated fitting procedures; however, recent progress suggests that these maps, too, have a high degree of regularity. Two boundaries are well defined by retinotopic features (sections 6.2.2, 6.2.3), and one of these also coincides with an anatomical landmark, the ptCoS. Moreover, the internal structure of the hV4 map and its neighbors are well understood, such that the iso-eccentricity and iso-angle lines derived from retinotopic mapping follow regular patterns.

### Variability across subjects

While the most of the large scale structures in the ventral occipital retinotopic maps are similar across subjects, there are also individual differences. Some of these differences likely reflect quantitative differences between subjects in the size and layout of the maps. Other differences reflect various sources of measurement noise. In no case will a measured map exactly match a template, and in some cases map boundaries will be ambiguous. We believe that the best approach offered is to simultaneously satisfy as many of the constraints from the anatomy, eccentricity, and angle maps as possible.

## ACKNOWLEDGMENTS

The authors acknowledge two funding sources, NIH Grant R00-EY022116 (JW) and NIH Grant R01-EY023915 to Kalanit Grill Spector (supporting NW).

### DISCLOSURES

The authors have nothing to disclose.

